## Supplementary material for "Gut microbiota DPP4-like enzymes are increased in type-2 diabetes and contribute to incretin inactivation": Table 1

Table 1. Estimation of EC_50_ in a gliptins dose-response assay.

|  | hsDPP4 | | | pmDPP4 | | |
| --- | --- | --- | --- | --- | --- | --- |
| Gliptin | EC_50_ | CI | R^2^ | EC_50_ | IC | R^2^ |
| Sitagliptin | 5.01 x10^-8^ | 4.76 x 10^-8^ to 5.27 x 10^-8^ | 0.995 | 9.04 x10^-5^ | 1.35 x 10^-5^ to 1 | ND |
| Vildaliptin | 1.54 x10^-7^ | 1.41 x 10^-7^ to 1.70 x 10^-7^ | 0.980 | 4.57 x10^-7^ | 3.68 x 10^-7^ to 5.72 x 10^-7^ | 0.833 |
| Saxagliptin | 4.29 x 10^-8^ | 4.14 x 10^-8^ to 4.44 x 10^-8^ | 0.998 | 1.45 x 10^-7^ | 1.30 x 10^-7^ to 1.69 x 10^-7^ | 0.971 |
| Linagliptin | 3.68 x 10^-9^ | 3.30 x 10^-9^ to 4.10 x 10^-9^ | 0.982 | 1.48 x 10^-5^ | 4.87 x10^-6^ to 1 | ND |

EC_50_, Estimated at molar (M) concentration. CI, confidence interval at 95%. R^2^, calculated as the fit of data into the non-linear function of the inhibitor concentration vs normalized response. Data obtained from independent experiments (N = 2). ND, data did not fit as non-linear function.
