## Additional files for "Gut microbiota DPP4-like enzymes are increased in type-2 diabetes and contribute to incretin inactivation"

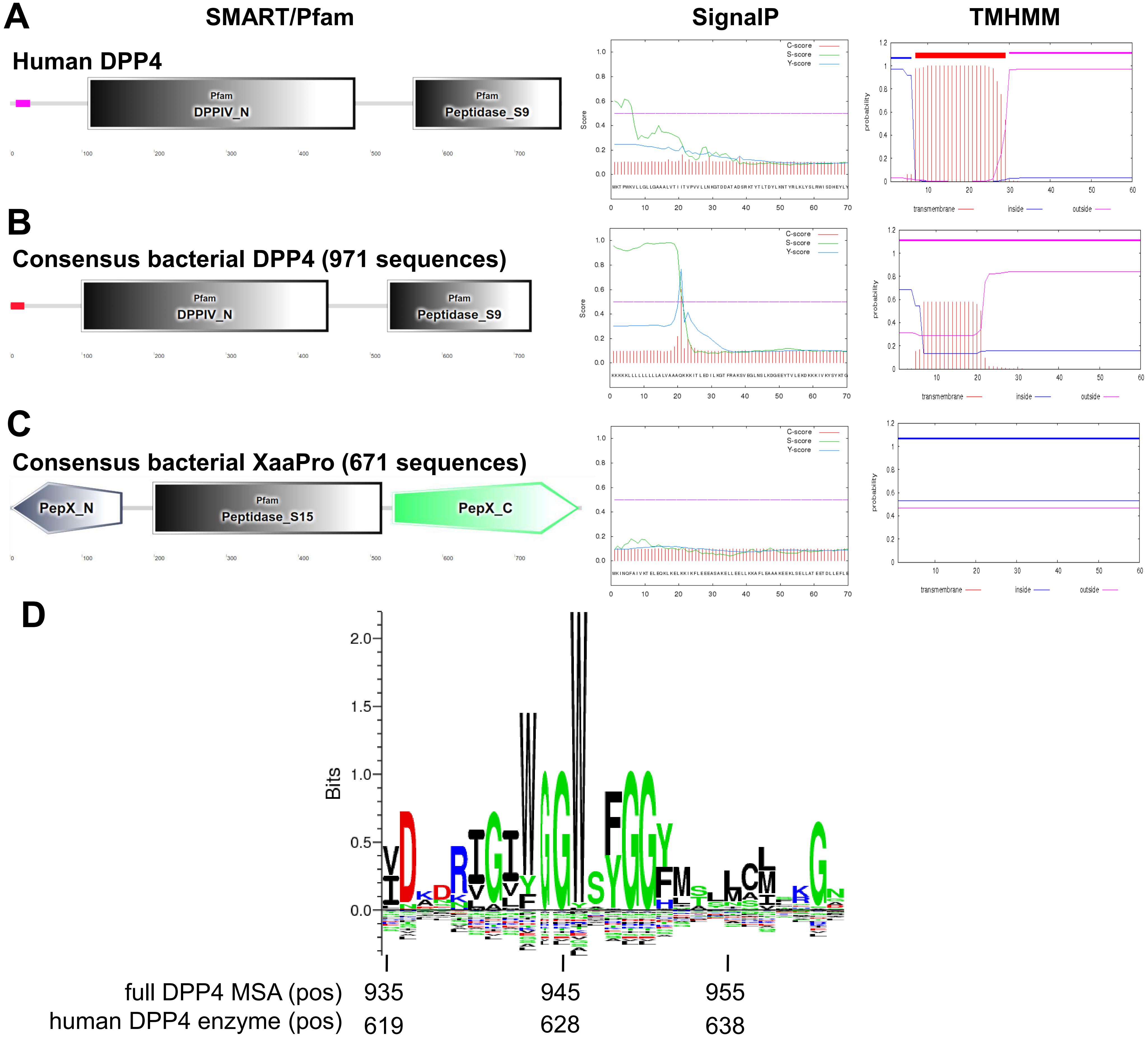


**Figure S1.** Comparison of DPP4-like function-associated peptidases. Domain architecture, signal peptide presence and transmembrane domain predictions were examined on the human DPP4 enzyme (SwissProt id P27487) (**A**), the consensus amino acid profile of bacterial DPP4-like proteins (**B**), and the consensus amino acid profile of PepX peptidases (**C**) using the respective web servers (top labels). Individual functional domains according to SMART and/or Pfam annotation present in full-length query proteins are presented in classical architecture diagrams. SignalP prediction for the N-terminal 70 amino acids shows C- (raw cleavage site score, red lines), S- (signal peptide score, green lines), and Y- (combined cleavage site score, blue lines) scores to determine the individual probability of amino acids to be part of a signal peptide driving protein secretion or potential membership of a transmembrane structure. TMHMM prediction for the N-terminal 70 amino acids shows the probability of individual amino acids to be part of alpha helices embedded in biological membranes, thus indicating localization in the transmembrane (red lines), inside of cell (blue line) or outside of cell (pink lines). The amino acid logo representation of the multiple sequence alignment (MSA) of DPP4-like proteins (bacteria, mouse, rat, and human) for the catalytic motif of S9 peptidase family is drawn in **D**. Amino acid logo was created using the positions between amino acids 935 and 941 of the MSA using the SeqLogo sever (http://www.cbs.dtu.dk/biotools/Seq2Logo/) and based on the P-Weighted Kullback-Leibler score. The amino acid positions for the human protein are also shown at the bottom. This regions shows the MEROPS [22] catalytic motif (GWSYGGY) of these serine proteases (S9).

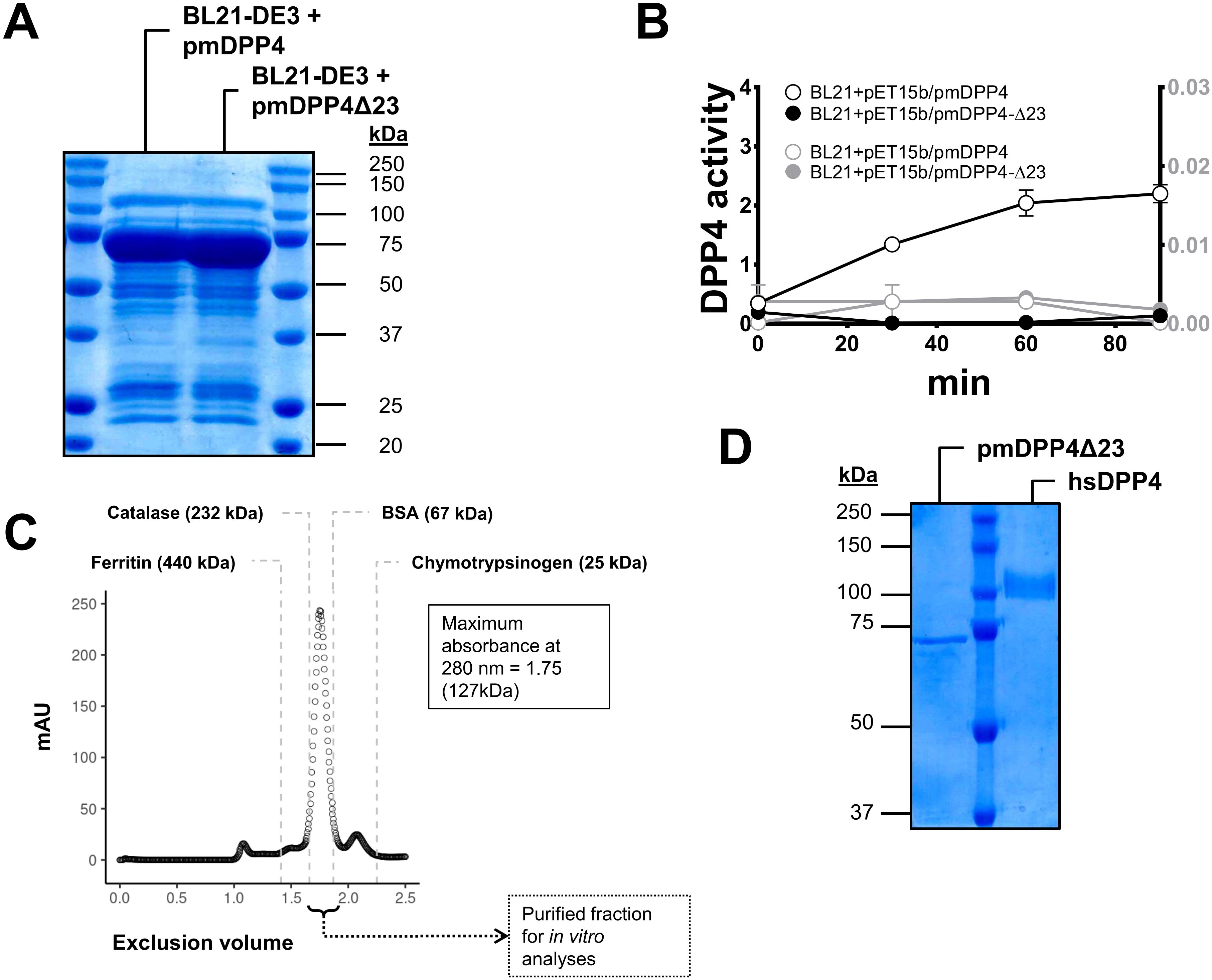


**Figure S2. A**, SDS-PAGE analysis of *E. coli* BL21-DE3 cell extracts expressing recombinant pmDPP4 and pmDPP4Δ23 proteins. **B**, Ninety minute kinetics of pmDPP4 activity (left y-axis, in terms of mU/mL units) and LDH activity (right y-axis, in terms of 490 nm - 680 nm absorbance) mediated by cell-free supernatants from *E. coli* BL21 cells carrying pET15b/pmDPP4 or pET15b/pmDPP4Δ23 constructs upon induction. **C**, Gel filtration assay of an ionic-exchange purified fraction of the pmDPP4Δ23 protein, revealing a dimeric nature in solution according to the calculated MW from the standards elution profile. **D**, Purified pmDPP4Δ23 and human DPP4 proteins used in the *in vitro* assays for human hormone hydrolysis and gliptin inhibition. Gels were stained with Coomassie blue to visualize protein bands. The Dual Xtra pre-stained protein standard (Bio-Rad, Hercules, CA) was used as a molecular size reference. LDH, lactate dehydrogenase.

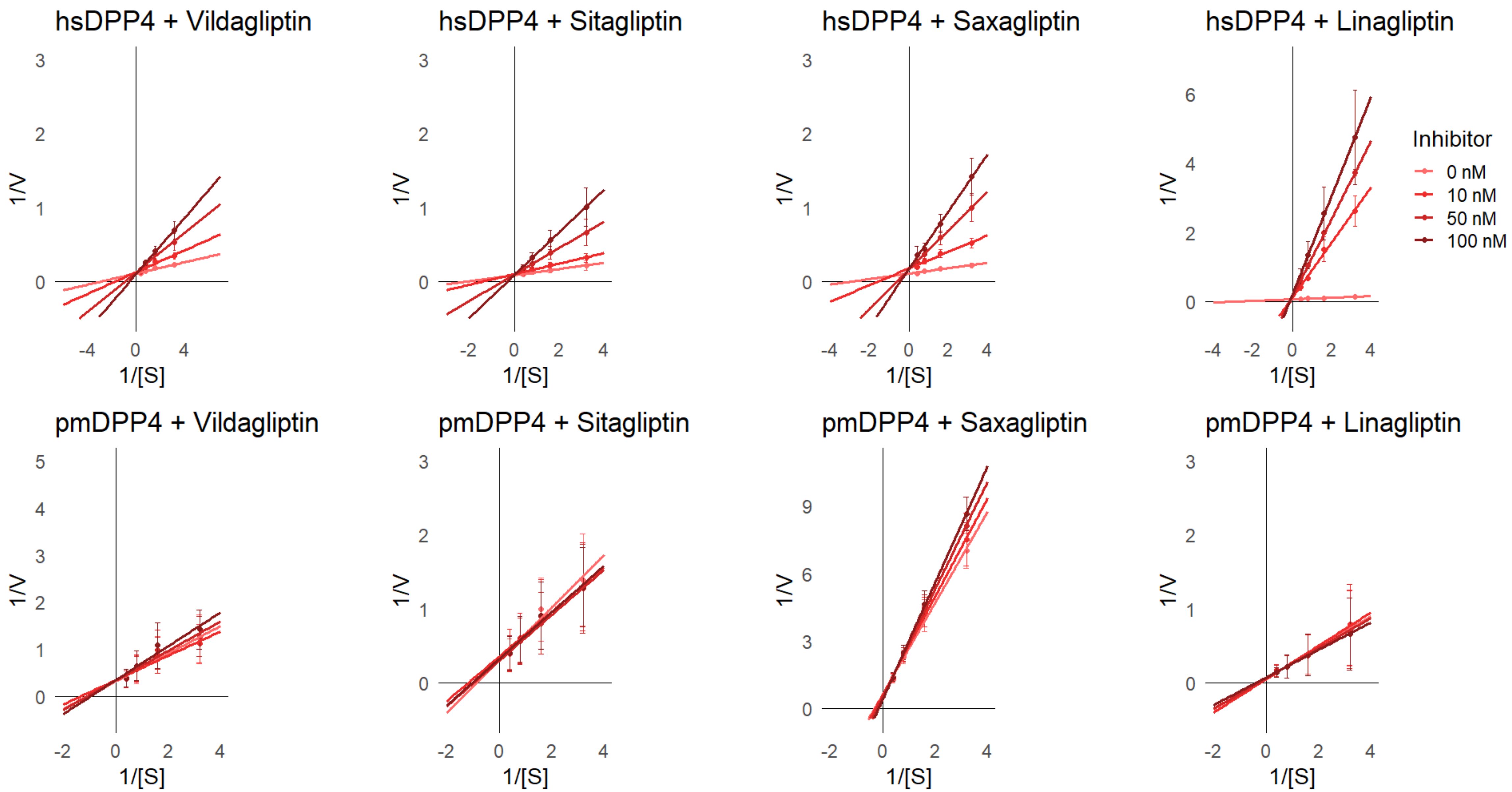


**Figure S3.** Gliptin inhibition kinetics on pmDPP4Δ23. Lineweaver-Burk double-reciprocal (1/V and 1/S) plots showing pmDPP4Δ23 (bottom panels) and hsDPP4 (top panels) gpPNA hydrolysis in presence of vildagliptin, sitagliptin, saxagliptin, and linagliptin. Inhibitor concentration is shown as line legend color from lower (0 nM, light red) to higher (100 nM, burgundy lines) concentrations. The apparent $K_m$ was calculated respectively from inhibitor-associated data points using linear regression to find intersection point at the y-axis (1/S). V, reaction velocity (nmol PNA $min^{-1}$ pmole $enzyme^{-1}$); S, substrate concentration (mM gpPNA).

| Enzyme | Inhibitor (nM) | Km (mM) | SD | Fold-change | P-value (t.Test) |
|---|---|---|---|---|---|
| *hsDPP4* | Vildagliptin 100 nM | 1.94 | 0.41 | 4.82 | 0.021 |
| *hsDPP4* | Vildagliptin 50 nM | 1.36 | 0.12 | 3.07 | 0.004 |
| *hsDPP4* | Vildagliptin 10 nM | 0.59 | 0.07 | 0.76 | 0.019 |
| *hsDPP4* | Vildagliptin 0 nM | 0.33 | 0.02 | 0.00 | |
| *hsDPP4* | Sitagliptin 100 nM | 4.35 | 0.60 | 12.58 | 0.007 |
| *hsDPP4* | Sitagliptin 50 nM | 2.15 | 0.27 | 5.73 | 0.007 |
| *hsDPP4* | Sitagliptin 10 nM | 0.72 | 0.03 | 1.24 | 0.000 |
| *hsDPP4* | Sitagliptin 0 nM | 0.32 | 0.03 | 0.00 | |
| *hsDPP4* | Saxagliptin 100 nM | 3.34 | 1.04 | 10.53 | 0.037 |
| *hsDPP4* | Saxagliptin 50 nM | 1.58 | 0.31 | 4.46 | 0.018 |
| *hsDPP4* | Saxagliptin 10 nM | 0.63 | 0.05 | 1.17 | 0.006 |
| *hsDPP4* | Saxagliptin 0 nM | 0.29 | 0.00 | 0.00 | |
| *hsDPP4* | Linagliptin 100 nM | 7.34 | 1.33 | 19.01 | 0.012 |
| *hsDPP4* | Linagliptin 50 nM | 8.24 | 1.10 | 21.48 | 0.006 |
| *hsDPP4* | Linagliptin 10 nM | 9.80 | 2.50 | 25.74 | 0.023 |
| *hsDPP4* | Linagliptin 0 nM | 0.37 | 0.05 | 0.00 | |
| *pmDPP4* | Vildagliptin 100 nM | 2.84 | 0.13 | 0.78 | 0.076 |
| *pmDPP4* | Vildagliptin 50 nM | 2.02 | 0.18 | 0.27 | 0.377 |
| *pmDPP4* | Vildagliptin 10 nM | 1.12 | 1.35 | -0.30 | 0.595 |
| *pmDPP4* | Vildagliptin 0 nM | 1.60 | 0.65 | 0.00 | |
| *pmDPP4* | Sitagliptin 100 nM | 1.51 | 0.15 | -0.15 | 0.301 |
| *pmDPP4* | Sitagliptin 50 nM | 1.44 | 0.16 | -0.19 | 0.214 |
| *pmDPP4* | Sitagliptin 10 nM | 1.25 | 0.17 | -0.30 | 0.096 |
| *pmDPP4* | Sitagliptin 0 nM | 1.79 | 0.34 | 0.00 | |
| *pmDPP4* | Saxagliptin 100 nM | 4.54 | 0.99 | 1.73 | 0.036 |
| *pmDPP4* | Saxagliptin 50 nM | 2.63 | 0.22 | 0.58 | 0.552 |
| *pmDPP4* | Saxagliptin 10 nM | 1.85 | 0.45 | 0.11 | 0.012 |
| *pmDPP4* | Saxagliptin 0 nM | 1.66 | 0.07 | 0.00 | |
| *pmDPP4* | Linagliptin 100 nM | 1.73 | 0.08 | 0.18 | 0.108 |
| *pmDPP4* | Linagliptin 50 nM | 1.66 | 0.11 | 0.13 | 0.196 |
| *pmDPP4* | Linagliptin 10 nM | 1.51 | 0.25 | 0.03 | 0.823 |
| *pmDPP4* | Linagliptin 0 nM | 1.46 | 0.18 | 0.00 | |

**Table S1.** $K_m$ fold-change on pmDPP4Δ23 and hsDPP4 enzymes upon gliptin inhibition. Data derived from kinetics assays shown in Figure S3 are shown in this spreadsheet. Headers indicate enzyme source, inhibitor tested, $K_m$ estimated, standard deviation (SD), fold-change, and p-value obtained from comparisons with respective reactions with no inhibitor.
